# Gauge-and-compass migration: inherited magnetic headings and signposts can adapt to changing geomagnetic landscapes

**DOI:** 10.1101/2022.06.29.498190

**Authors:** James D. McLaren, Heiko Schmaljohann, Bernd Blasius

## Abstract

**Background:** For many migratory species, inexperienced (naïve) individuals reach remote nonbreeding areas independently using one or more inherited compass headings and, potentially, magnetic signposts to gauge where to switch between compass headings. Inherited magnetic-based migratory orientation programs have not yet been assessed as a population-level process, particularly where strong geomagnetic spatial gradients or long-term shifts could create mismatches with inherited magnetic headings. In particular, it remains unstudied whether and how, under natural selection, inherited headings and signposts could potentially adapt to long-term geomagnetic shifts.

**Methods:** To address these unknowns, we modelled bird migration using an evolutionary algorithm incorporating global geomagnetic data (1900-2023). Modelled population mixing incorporated both natal dispersal and trans-generational inheritance of magnetic headings and signposts, the latter including intrinsic (stochastic) variability. Using the model, we assessed robustness of signposted and non-signposted trans-hemispheric songbird migration across a rapidly magnetically-shifting Nearctic breeding region (mean 34° declination shift) via Europe to Africa.

**Results:** Model-evolved magnetic-signposted migration was (i) overall successful throughout the 124-year period, with 60-90% mean successful arrival across a broad range in plausible compass precision, (ii) through reduced trans-Atlantic flight distances, up to twice as successful compared with non-signposted migration, but (iii) to avoid evolving unsustainable open-ocean flights, intrinsic variability in inheritance of magnetic headings was required (model-evolved σ ≈ 2.6° standard error in inherited headings).

**Conclusions:** Our study supports the potential long-term viability of inherited magnetic migratory headings and signposts, and illustrates more generally how inherited migratory programs can both mediate and constrain evolution of routes, in response to global environmental change.

## Introduction

Myriads of migrating animals undertake seasonal journeys across regional to cross-continental-scales (1,2). For many migratory populations, seasonal routes are primarily mediated culturally, i.e., by collective and social cues (3,4). However, many long-distance migrants such as butterflies, sea turtles and night-migratory songbirds, migrate largely independently (5–7). Experienced solo-migrants can develop a map sense to navigate (reach known destinations from unfamiliar locations), for example by extrapolating magnetic field components (6,8,9). Inexperienced (hereafter, naïve) solo-migrants are thought to rely strongly on endogenous programs mediated by circannual timing and inherited compass headings (10–13). In the simplest case, a “clock-and-compass” migrant following a single inherited migratory heading would, depending on its primary migratory compass, result in either a geographic (e.g., star) or magnetic compass course, or else a gradually-shifting sun compass course (14–16). Many migration routes are indeed potentially explainable by one or more compass courses (14,15), contingent upon possessing sufficient compass precision (16,17) and ability to negotiate currents (18,19). However, many other migratory routes require distinct direction changes (often termed *Zugknicks* in bird migration), e.g., to avoid ecological barriers (20,21), or to exploit favourable habitats (22,23) or supportive current systems (9,24).

The mechanisms underlying how naïve migrants reliably mediate critical direction changes along unfamiliar routes remain unclear. Purely clock-mediated direction changes could prove unreliable given inherent variability in migratory schedules (25,26). Alternatively, naïve migrants could potentially take advantage of the broad-scale latitudinal structure of the Earth’s magnetic field, with both geomagnetic inclination (the vertical tilt of the geomagnetic field) and total field intensity decreasing from high to low latitudes (10,27). Experimental evidence suggests that naïve oceanic (9) and avian (10,28) migrants indeed can use geomagnetic information to mediate orientation shifts, but is inconclusive regarding which magnetic components are used and the extent to which updated headings are either extrapolated *in situ* or predetermined, i.e., inherited. If the former, naïve migrants have been proposed to reach inherited magnetic “signature” locations by following perceived or extrapolated gradients in bi-coordinate geomagnetic components, e.g., inclination and intensity, (9,29). Alternatively, naïve migrants could potentially mediate switches (*Zugknicks*) between a fixed sequence of inherited headings upon passing inherited magnetic signposts, e.g., a threshold value of inclination or intensity (6,9,10). In this study, we assess such a “gauge-and-compass” migratory orientation program, which does not require migrants to reach a specific geomagnetic location, nor to extrapolate between experienced geomagnetic configurations.

A critical factor affecting feasibility of migration based on inherited magnetic information is its robustness to spatiotemporal geomagnetic variability, which could create mismatches with inherited magnetic headings. The Earth’s magnetic field is irregularly aligned with the true geographic N-S axis, according to geomagnetic declination (clockwise angle from true to magnetic N), and undergoes temporal fluctuations between daily and centuries-long time scales (30). Both spatial and temporal geomagnetic changes can occur rapidly in some areas such as East-arctic Canada and Greenland (Fig. 1a), where the N magnetic pole is drifting an order of magnitude faster than a century ago (31), causing extreme declination shifts (Fig. 1b). Therefore, for migratory birds, it is often thought that orientation or navigation through strongly varying geomagnetic fields is intractable without regular calibration using non-magnetic cues (32–34). However, it is also underappreciated that spatial geomagnetic gradients could enhance the feasibility and efficiency of goal-directed movement (35). For example, with an Eastward increase in declination (in the Northern Hemisphere), Southward geomagnetic headings will partially correct for erroneous displacement, e.g., by wind (Suppl. Fig. 1).

**Figure 1.**
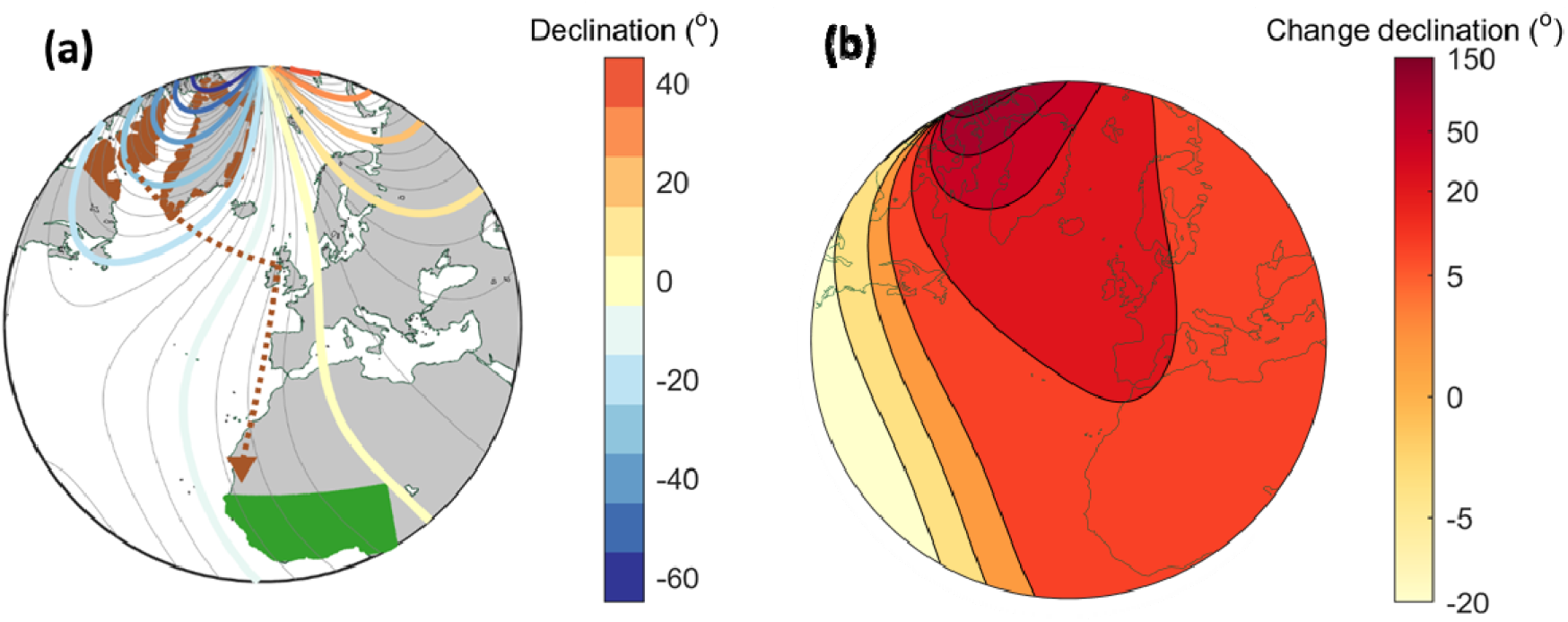
Extreme geomagnetic-field gradients and temporal shifts along migration routes of northern wheatears (*Oenanthe oenanthe leucorhoa*). (**a**) Contours of geomagnetic declination in 2010 (degrees clockwise from true to magnetic North; colour scale on right), with modelled natal range (in brown) and wintering grounds (goal area; green). The brown dashed line and arrow depicts the approximate actual route from Iqaluit, Baffin Island taken by a juvenile *leucoroha* northern wheatear tagged in 2010 using light-level geolocation (44). Distance-minimizing great circle routes are straight lines in the stereographic projection (35); (**b**) Contours of (clockwise) changes in declination from 1900 to 2023, with colour scale on right.

Regarding robustness to temporal variability, it is important to consider whether plasticity in inheritance of orientation cues (11,36,37) can track temporal geomagnetic shifts at a population level. In order to assess this, since migratory headings will vary geographically (16,37,38), and are typically inherited as averages (11,39, but see 38), it is important to account for trans-generational changes (mixing) in orientation (11,37). For example, natal dispersal will create population-level variation among individual migratory headings (40,41). More generally, intrinsic stochastic variability in inheritance of traits - as distinct from population-level variation - sometimes referred to as bet-hedging, is expected to be beneficial in unpredictably varying environments (36,42,43). To assess the long-term viability of inherited magnetic-based migration at the population level, including the benefit of inherited magnetic signposts, we developed a model of migration through spatiotemporally-varying geomagnetic landscapes using an evolutionary algorithm approach (45,46). The migration model is based on compass-based movement (16), extended to include signposted directional switches in the migratory direction. The evolutionary algorithm mimics inheritance of migratory orientation among successful migrants, accounting for both spatial population mixing (through natal dispersal) and stochastic variation in inheritance (36,43,47). Being specifically interested in inherited migratory orientation rather than population dynamics, we did not vary the population size or breeding locations, and considered only naïve migrants, by repopulating new modelled “offspring” each year based on natal dispersal of successful migrants from the previous year. The model first “evolves” a viable test population for the migration route considered, i.e., one adapted to both geomagnetic data from random years and the initial test year (1900), then tests its robustness to 124 years (1900-2023) of global geomagnetic change (48). During optimisation of the viable migratory population, the model additionally “evolves” the magnitudes of intrinsic variability (standard variations in inheritance) in inheritance of headings, signposts and natal dispersal. Contrastingly, when assessing robustness of the viable population, we more conservatively kept intrinsic variability of these three quantities fixed.

Migratory flight can involve magnetic headings in up to three ways: through inheritance of magnetic as opposed to geographic headings, or through use of either a primary or an in-flight migratory compass (33,49). With inheritance of geographic compass headings, the magnetic compass headings can still be imprinted, e.g., at the natal site (49). With a primary magnetic migratory compass, flight directions on departure (e.g., nightly) are determined relative to the proximate geomagnetic field. An in-flight magnetic compass can also be cue-transferred on departures from a primary geographic (star) or sun compass (16,33). To focus on magnetic-based migration, we by default modelled inherited magnetic headings with both a primary and in-flight magnetic compass. For comparison, we also tested each combination of geographic and magnetic compass involvement (as inherited, primary and in-flight compass).

We considered three inherited migratory orientation programs for naïve bird migrants: non-signposted migration, i.e., following a single inherited heading, and two signposted migratory programs, based on either geomagnetic inclination or intensity. With an inherited signpost, modelled migrants shift to a second inherited (*Zugknick*) compass heading once the perceived magnitude in inclination or intensity falls below an inherited threshold value. Using the viable population as a starting point, each migratory orientation program was assessed by arrival success over the 124-year simulated period. As a sensitivity analysis, we also assessed the effect of migrants’ precision, both in sensing the magnetic field components and in overall flight-step direction (16,50). As an uncertainty analysis, we also tested declination-signposted migration, which would require extrapolation between a geomagnetic and geographic reference (e.g., via a star or sun compass).

We chose to model a migratory songbird, the East-Nearctic-breeding population of the northern wheatear (*Oenanthe oenanthe leucorhoa*, hereafter *leucorhoa* wheatear). This subspecies faces a clear energetic and survival bottleneck, the Atlantic Ocean, *en route* to their wintering grounds in sub-Sahelian West Africa (20,51), while also traversing strong geomagnetic gradients in a rapidly-shifting polar geomagnetic landscape (Fig. 1, 31). *Leucorhoa* wheatears can potentially reach Spain or even Africa in several days of non-stop flight (51,52) but are known to detour (*Zugknick*) via North-West Europe, with tracked migrations (Fig. 1a) from Baffin Island via Britain or Ireland (44) and from Southwest Greenland via France and Spain (52,53). Long-distance bird migrants typically intersperse periods of daily or nightly flight-steps with extended periods of stopover, particularly where energy reserves are readily acquired and most needed (64,65). To prepare for their trans-Atlantic flights, *leucorhoa* wheatears are known to accumulate massive energy reserves in autumn (up to double their body mass, 53,54). They are otherwise open habitat generalists and pass across Europe on a broad-front (55,56). For simplicity and to focus on orientation and geomagnetic effects, we modelled flight energetics as potential flight hours, replenished during extended stopover periods. Similarly, to focus on inheritance of migratory orientation rather than population demographics, we assessed individual fitness by only successful arrival within their modelled wintering grounds (44,56).

Our study represents a strict test of both magnetic-based inheritance and signposted migration under geomagnetic change. Given the mortality risk over the ocean barrier, we predict that signposted migration benefits successful arrival of *leucoroha* wheatears in Africa. Furthermore, given the West-East gradient in declination (Fig. 1a, Suppl. Fig. 1), we expect that primary magnetic headings will be more successful compared with geographic (e.g., star compass) headings. Finally, we predict that inheritance of magnetic orientation benefits from intrinsic variability (42,43) beyond natal dispersal and population mixing. More generally, our study highlights how natural selection might enable migratory populations to adapt to global changes in a key environmental migration cue.

## Methods

### Circular quantities and precision

Migratory headings were assumed to be inherited as clockwise angles relative to proximate geographic or magnetic N-S, according to the inherited compass. Angular quantities, i.e., headings and inclination or declination signposts, were sampled using a von Mises distribution, the circular equivalent of a normal distribution, governed by the von Mises concentration, κ (57). For interpretability, we report circular precision and variability by 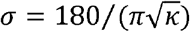, nearly equivalent to circular standard deviation for σ < 30° (16,57).

### Evolutionary strategy algorithm

#### Overview

Evolutionary algorithms originated to optimise logistical and workflow problems, through mimicking the iterative process of natural selection (46). In particular, intrinsic variability in inheritance of traits, e.g. through concurrently “evolving” a standard deviation in inheritance, can accelerate convergence to optimal solutions (43,46). Many classes and variations of evolutionary algorithms have been developed, including to answer biological questions (45,46). One such class, evolutionary strategy algorithms, combines natural selection and inheritance of traits as real-valued parameters, incorporating both mutations and recombination in subsequent model iterations (43,45).

We developed an evolutionary strategy algorithm to model (micro-)evolution of inherited migratory headings and signposts, based on successful arrival of naïve (first-fall juvenile) migrants to their wintering ground, and on subsequent population mixing. Table 1 lists the range of default and extended ranges of key model parameters. As mentioned, we only retained next-generation (offspring) naïve migrants in the simulation; this is a common technique in evolutionary strategies (known as a comma-strategy as opposed to a plus-strategy, 43,58). Population mixing depended on two coupled model-evolved processes: natal dispersal, and inheritance of headings and signposts. Specifically, two successful “parents” were randomly selected for each departure location with a probability weighted by the resulting natal dispersal distance (40,59). Parental migratory traits were passed on to the next generation as an average (11,37), together with intrinsic (stochastic) variability (36,60), as outlined below.

**Table 1.** Key model parameters and their default values, regarding migration and inheritance, for assessment of orientation programs.

| Quantity | Default | Sampled | Distribution | Fixed or evolved (in full simulation) | Sensitivity or uncertainty analysis |
| --- | --- | --- | --- | --- | --- |
| Population size | 50,000 | - | - | fixed | 1000 – 250,000 |
| Duration of spin-up simulation | 75 years | 50 random years, 25 years for 1900 | uniform | - | 25 – 150 years |
| Duration of full simulation | 124 years (1900 – 2023) | yearly | - | - | unchanging field (1900) backwards (from 2023) |
| Natal locations | in viable habitat | initially, for spin-up | uniform | fixed | - |
| Flight capacity | 48 – 72 h | initially, replenished per stopover | uniform | fixed | - |
| Primary compass | magnetic | per flight-step | circular | fixed | star compass |
| In-flight (nightly) headings | magnetic | per flight-step | circular | fixed | star compass |
| Flight-step precision | $\sigma = 15^\circ$<br>($\kappa = 14.6$) | per flight-step | circular | fixed | $5^\circ - 45^\circ$<br>($\kappa = 1.6 - 131.3$) |
| Precision gauging inclination | $\sigma = 5^\circ$<br>($\kappa = 131.3$) | in-flight hourly | circular | fixed | $0.1^\circ - 20^\circ$<br>($\kappa = 8 - 3.3 \cdot 10^5$) |
| Precision gauging intensity | $\sigma = 2\%$ | in-flight hourly | normal | fixed | 0.1 – 20% |
| Natal dispersal (expected mean) | 25 m – 25 km | from model spin-up | uniform (initial spin up) | fixed | 10 m – 250 km (fixed) |
| Natal dispersal (realized) | derived from distribution | yearly among successful migrants | half-normal, averaged between parents | fixed | - |
| Inherited headings | magnetic | yearly among successful migrants | circular, averaged between parents | evolved | geographic |
| Variability in inherited headings | $\sigma = 0.025^\circ - 5^\circ$<br>($\kappa = 131 - 5.3 \cdot 10^6$) | population mean from model spin-up | uniform (initial spin up) | fixed | - |
| Inherited signposts | geomagnetic inclination or intensity | yearly among successful migrants | uniform (initial spin up) | evolved | geomagnetic declination |
| Variability in inheritance of inclination signposts | $\sigma = 0.025^\circ - 1^\circ$<br>($\kappa = 131 - 3280$) | population mean from model spin-up | uniform (initial spin up) | fixed | - |
| Variability in inheritance of intensity signposts | $\sigma = 0.1 - 1\%$ | population mean from model spin-up | uniform (initial spin up) | fixed | - |
| Variability in inheritance of natal dispersal | 2.5 m – 1 km | during model spin-up | uniform (initial spin up) | none in full simulation | - |

#### Model spin-up and viable population

To derive a viable initial migratory population, analogously to with model spin-up in climate models (18,61,62), we first evolved modelled inherited headings, signposts and natal dispersal using geomagnetic data from random years and the initial year (1900). This was done in two phases: for each year of the 50-year first phase, geomagnetic data from a random year was used. Then, in a second phase, to allow inherited traits to adapt to the first modelled year, 25 years were simulated using geomagnetic data from 1900. For computational efficiency in convergence, during the first phase, we additionally (i) “evolved” the extent of intrinsic variability (standard deviation) in inheritance of three traits: headings, signposts and expected mean natal dispersal, at an individual level (36,43); (ii) in addition to natal dispersal, weighted selection of the second candidate parental migrant by its similarity in variability to the first selected parent (equivalent to assortative mating (45), i.e., with zero weight when maximally different and a maximal weight of one when identical). For the second spin-up phase and resultant viable population, intrinsic variability of headings and signposts were conservatively set to their (evolved) population-mean values. This is equivalent to assuming that, within the evolutionarily short (124-year) period considered, microevolution of intrinsic variability in inheritance may be constrained, e.g., by limits in plasticity through gene replication and expression (60,63).

#### Natal dispersal

For the return to the breeding area, we modelled natal dispersal as a half-normal probability distribution with distance (59). For a given natal location, the probability of selecting a candidate successful migrant as “parent” depends on the ratio, *r*, of its resultant dispersal distance *d* to the (expected) mean dispersal, *D_N_* through a (Gaussian) exponential *p*(*r*)~*e*^-*r*_2_/π^(Fig. 2). Therefore, the relative selection probability among successful individuals with ratios r_i_ is readily computed as 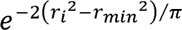, where *r_min_* is the minimum value in *r* among successful migrants from the previous year. During the first (50-year) spin-up phase, the model evolved the expected mean dispersal, *D_N_*, as an individual trait (initially uniformly distributed between 25-m and 25-km) and also the intrinsic variability (standard deviation) in inheritance of mean dispersal (initially uniformly distributed between 2.5-m and 1-km). For the second spin-up phase and resulting viable population, although natal dispersal may be an inheritable trait (40,41), we chose a more conservative approach by setting to its evolved population mean, as determined in the first spin-up phase, without intrinsic variability. As a sensitivity analysis, we also simulated migration with population-uniform (non-evolved) distributions of natal dispersal with expected means ranging from 10-m to 250-km.

**Figure 2:**
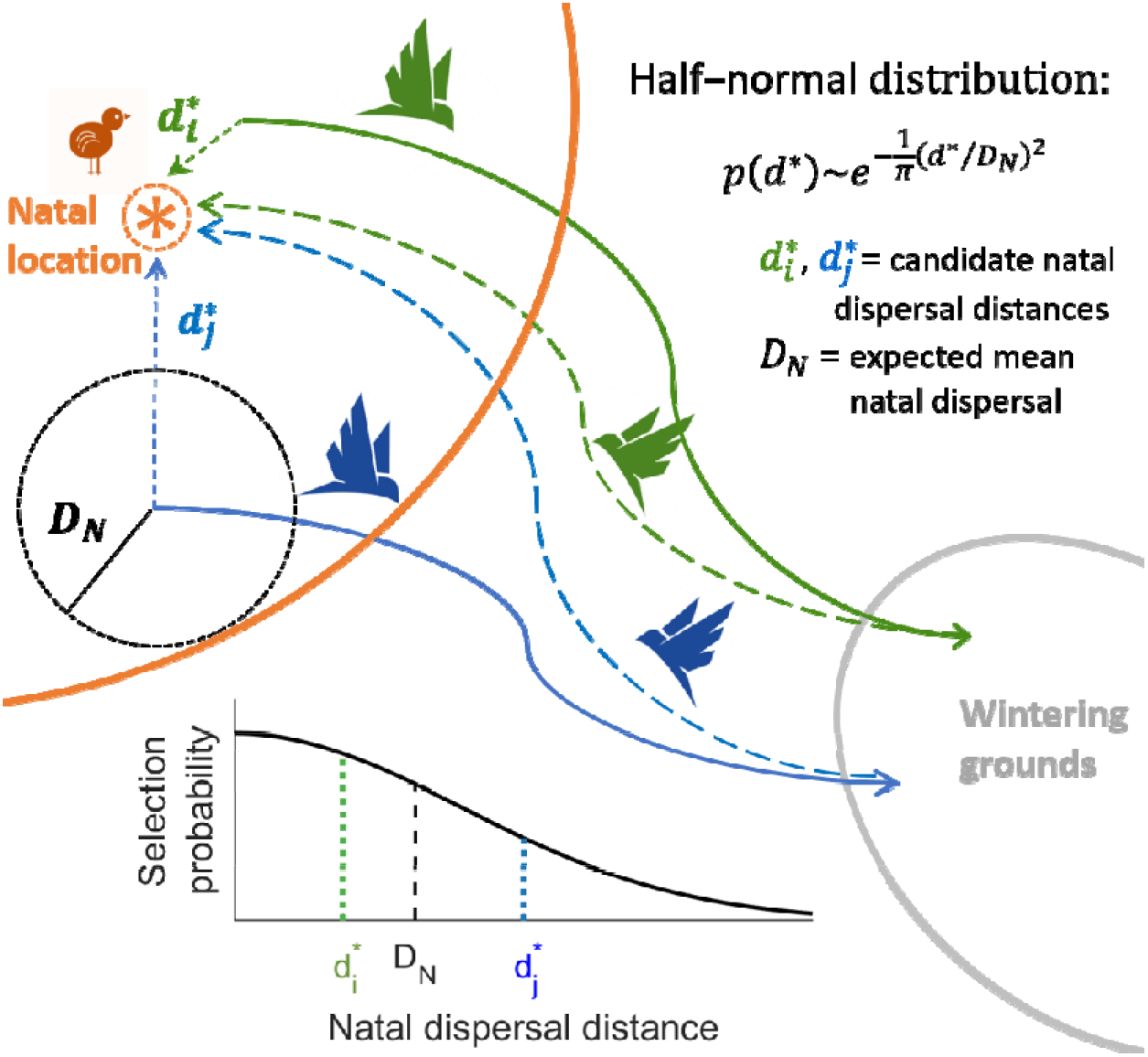
Model selection of parental migrants based on natal dispersal. For each breeding location in each modelled generation (year), selection among candidate successful migrants from the previous year is based on a half-normal distribution (lower left) given the resultant dispersal distance, *d*, and expected mean natal dispersal, *D_N_* (equation, top right). During the initial (50-year) model “spin-up”, to create a viable migratory modelled population, the expected mean, *D_N_*, was “evolved” as an individual trait (see text). For the actual assessment of robustness to geomagnetic change (1900-2023), natal dispersal was set uniformly with expected mean set to the population mean value from the model spin-up, i.e., not treated as “inheritable” or “evolvable”.

### Wheatear migration model

#### Overview

We modelled naïve *leucorhoa* wheatears migrating from natal areas in Greenland and North-East Canada (10°-80°W and 57.5°-84°N) to their wintering grounds (hereafter, goal area) in sub-Sahelian West Africa (20°W - 5°E and 5°-15°N). To focus on robustness of inherited magnetic-based orientation rather than feasibility of *leucoroha* wheatear migration *per se*, the model rules were designed to provide both realistic and potentially sufficient compass precision and energy reserves to arrive at the goal area. We first outline the modelled migratory process, then describe the initialization of model parameters, model implementation and model assessments.

#### Natal locations and initial departure

Natal locations were held fixed throughout each simulation. These were initially set to random locations with at least 10% low vegetation and less than 15% barren habitat within a 1°x1° area, based on EarthEnv Global 1-km Consensus Land Cover (66), producing a distribution closely resembling the known breeding range (cf. Fig. 1a with 56,67). We assumed migrants departed on average on August 20^th^ with a 5-day standard deviation, but between August 6^th^ and September 3^rd^ (44,56).

#### Flight steps and identification of signposts

Modelled birds flew at constant 15 m/s ground speeds (accounting for a mean tailwind, 51,54), following either constant magnetic or geographic (star) compass headings, which were updated hourly. Flight-steps lasted from 90 minutes after sunset until 90 minutes before sunrise (54), for minimally 6 hours and maximally 12 hours, or until land was in sight. Flight durations were calculated using the algebraic formula for sunset hour for a spherical Earth based on date and latitude (68, and see 16). We considered a 15° default precision among flight-steps (κ=14.6), consistent with in-flight measurements of migrating songbirds (69,70) and model predictions of required precision (16,19). With signposted migration, individuals switched headings once on land and when their perceived inclination or intensity fell below the inherited (threshold) magnitude. We assumed conservatively that, to identify signposts, migrants could gauge magnitudes of inclination within 5° (71) and within 2% of (IGRF) field intensity (i.e., ca. 5000 nT at mid-latitudes (49,50)). For sensitivity analysis, we assessed migration with 5°-45° directional precision among flight-steps, 0.1°-20° precision in gauging inclination, and 0.1-20% in intensity.

#### Energy reserves and stopovers

Given the initial migratory endurance flight, modelled flight capacities were set to uniformly randomly sampled to between 48-72 hours, roughly equivalent to 62-106% relative gain in body mass as fat (64), as regularly observed among *leucoroha* wheatears in the wild (53,54). Extended migratory stopovers, assumed to last 5 days, occurred whenever potential flight ranges fell below a threshold (set to three nightly flight durations), or following flight-steps which began at sea (21). For simplicity and feasibility to cross the Mediterranean Sea and Sahara desert (18), energy reserves (mainly fat) were replenished at stopovers for 48-72 hours of potential flight (sampled uniformly, but minimally as per on arrival). Refuelling was however not permitted in barren land (based on 0% vegetation or NDVI within 1°x1° area, 66). If still over water at dawn, modelled individuals stopped at the nearest viewable coastal point (on a ca. 20-km grid), which was identified at each of the last 3 deciles of the nightly flight, based on a detection probability which decreased linearly with increasing distance up to 300 km (i.e., land immediately on the coast was always detectable, to 50% of the time at 150 km, to never beyond 300 km). If no land was viewable, modelled migrants flew until the next dusk, stopping at the nearest viewable coastal point at each decile of flight, or flying until energy reserves were depleted (mortality).

#### Arrival success

Migrants were considered successful if they arrived in the modelled goal area within a default of 90 days after leaving the breeding area (44,56). Signposted migration was still considered successful if migrants arrived in the goal area without having detected a signpost (e.g., when the magnitude of the inherited signpost fell below that of the relevant geomagnetic field component at the arrival location). However, individuals were considered unsuccessful if they overshot the goal area beyond half its latitudinal or longitudinal width (here, 15° in longitude or 5° in latitude), or flew poleward beyond 87.5°N.

#### Initialising orientation traits and variability

In the first (spin-up) year, initial inherited headings were set to a population mean value of the average of loxodrome (rhumbline) and great-circle headings between each departure location and randomly sampled candidate arrival locations within the goal area (wintering grounds), plus a circular random offset (the candidate arrival locations were not further used during the simulations). The random offset (κ = 0.033) was chosen such that its equivalent standard deviation (~17.5°) covered 75% of differences between all candidate great-circle and loxodrome headings. Initial values for second (signposted) inherited headings were permutated randomly from this pool. Initial values for inherited signposts were set to a random geomagnetic value between those of the departure and candidate arrival locations in the goal area, plus a noise term based on the precision in gauging signposts. During the first (50-year) spin-up phase, the model evolved the magnitudes of intrinsic variability (standard deviations) in inheritance of headings (initially sampled between 0.025°-5°), inclination and declination signposts (sampled between 0.1°-1.0°) and intensity signposts (between 0.1-1%).

### Model implementation

The model was implemented in MATLAB using the parallel programming, statistics and mapping toolboxes, as well the external Climate Data Toolbox (72) and IGRF data (48), the latter using a package adapted for parallel processing (73), updated for the most recent period (2015-2025, 74). Computation of topographic (coastline) and geomagnetic cues was calculated for all individuals in parallel, as was yearly population mixing (natal dispersal, parent-selection and trait inheritance).

Regarding computation time, a 75-year spin-up and 124-year simulation with 50,000 modelled individuals took approximately 5 hours to run for hourly-updated magnetic compass in-flight headings on a laptop with an 10^th^ generation Xeon ^©^ Intel chip (~3 hours for star compass in-flight headings).

### Model assessment

We assessed each migratory orientation program by its long-term geometric mean in yearly arrival success, 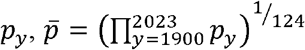. Geometric as opposed to arithmetic means are most appropriate for fitness or survival assessments, through accounting for negative effects of infrequent low success (36,42). We also compared evolution of *Zugknick* locations and kept track of mortality over water. To confirm that the default population size (50,000) produced reliable results (evolved orientation and resultant arrival success), we replicated intensity-signposted migration six times for seven different population sizes (between 100 and 250,000). We further confirmed that model results reflected effects of long-term geomagnetic shifts (e.g., as opposed to lack of convergence in viable orientation) by comparing long-term trends in arrival success to when simulating the 124-year period with geomagnetic data either from a single season (1900) or in reverse chronological order (from 2023 to 1900). Lastly, to test the benefit of intrinsic variability in inheritance of orientation traits, we simulated migration with perfect inheritance (averaging) of parental headings and signposts.

## Results

Model-evolved signposted *leucorhoa* wheatear migration to West Africa was overall and consistently successful across modelled years (1900-2023), with signposted detours over Europe resulting in higher arrival success compared with non-signposted migration. Fig. 3 depicts sample modelled trajectories from 2023 for each migratory program based on default model parameters (Table 1). With non-signposted migration (Fig. 3a), less than half of the individuals arrived successfully (43.5±1.2% among years), with frequent over-water mortality (53.8±0.8%) except for shorter ocean crossings such as from Northeast Greenland, or where trajectories came within sight of the Azores, permitting a stopover. With inclination-signposted migration (Fig. 3b), arrival success was higher (67.1±1.1%) and mortality over water more moderate (28.8±1.3%). Intensity-signposted migration (Fig. 3c) almost completely avoided the longest ocean-crossings, resulting in the highest arrival success (79.8±1.5%) and lowest over-water mortality (16.7±1.1%). Nonetheless, success dropped off slightly between 1900-2023 with intensity-signposted (2.6%) and inclination-signposted (1.6%) migration, but increased slightly with non-signposted migration (2.7%). With geomagnetic data parameterized in reverse chronological order (2023-1900; dashed lines in Fig. 1d), modelled arrival success was overall slightly (1-2%) higher for all programs, with the differences in success in 2023 and 1900 also reversed (6.1% increase for intensity-signposted, 4.5% increase for inclination-signposted, and 2.5% decrease for non-signposted migration), consistent with success being driven by geomagnetic effects. Using modelled geomagnetic data from 1900 for each year, arrival success was ~5% higher for signposted migration but ~5% lower for non-signposted migration. Results were robust to modelled population size with at least half the default number (25,000 or more) individuals, and also replicable regarding arrival success, mean inherited headings and *Zugknick* latitudes (Suppl. Fig. 2). With 10,000 or less individuals, between-replicate variability more than doubled regarding arrival success (Suppl. Fig. 2a) and evolved headings and *Zugknicks* (Suppl. Fig. 2a), with success decreasing significantly with 1000 or less modelled individuals.

**Figure 3:**
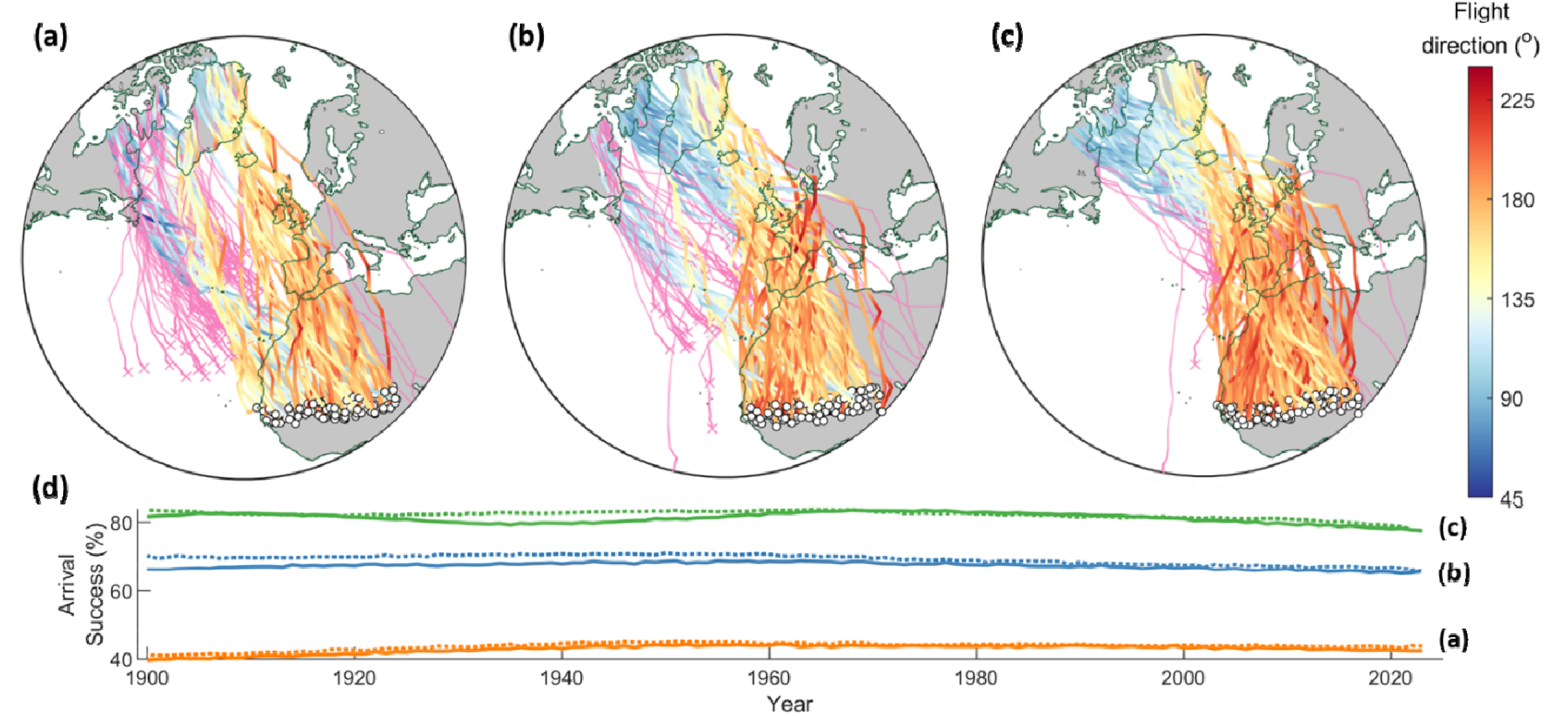
Model-evolved migratory trajectories and arrival success of inexperienced (naïve) *leucorhoa* wheatears (see Fig. 1). Random**ly**-sampled predicted trajectories from 2023, colour-coded to flight direction (degrees clockwise from geographic N) based on (**a**) non-signposted mig**rat**ion, following a constant inherited magnetic heading, (**b**) a magnetic signpost based on inclination and (**c**) a signpost based on geomagnetic intens**ity.** The results in (a)-(c) are with default model parameters (see Method and Table 1). For (**b**) and (**c**), encountering a signpost (inherited threshold geo**ma**gnetic value) triggers a shift to a second model-evolved inherited heading. Successful arrival in Africa is indicated by white circles, and pink tracks r**epr**esent unsuccessful individuals. Straight lines represent great circle routes in the stereographic azimuthal projection; (**d**) Arrival success (percentage **of** population) for non-signposted (solid orange line), intensity-signposted (solid blue line) and intensity-signposted migration (solid green line). Dash**ed** lines depict success when the model is parameterised by geomagnetic data in reverse chronological order (2023 to 1900).

For all three migratory orientation programs, arrival success depended strongly on precision among flight-steps (Fig. 4a). While the hierarchy between migratory programs remained consistent (intensity-signposted > inclination-signposted > non-signposted), the difference among them also decreased with lower flight-step precision. Contrastingly, all three programs were relatively robust to the degree of precision in gauging inclination and intensity to identify signposts (Fig. 4b). Similarly, each orientation program was relatively robust to the width of (non-evolved) distributions of natal dispersal, for magnitudes of expected mean dispersal up to 250 km (Fig. 4c). Model-evolved means in natal dispersal (hexagons, Fig. 4c) were ~16 km for each default orientation program (as per Fig. 3), resulting in arrival success close to the highest among fixed-distribution simulations.

**Figure 4.**
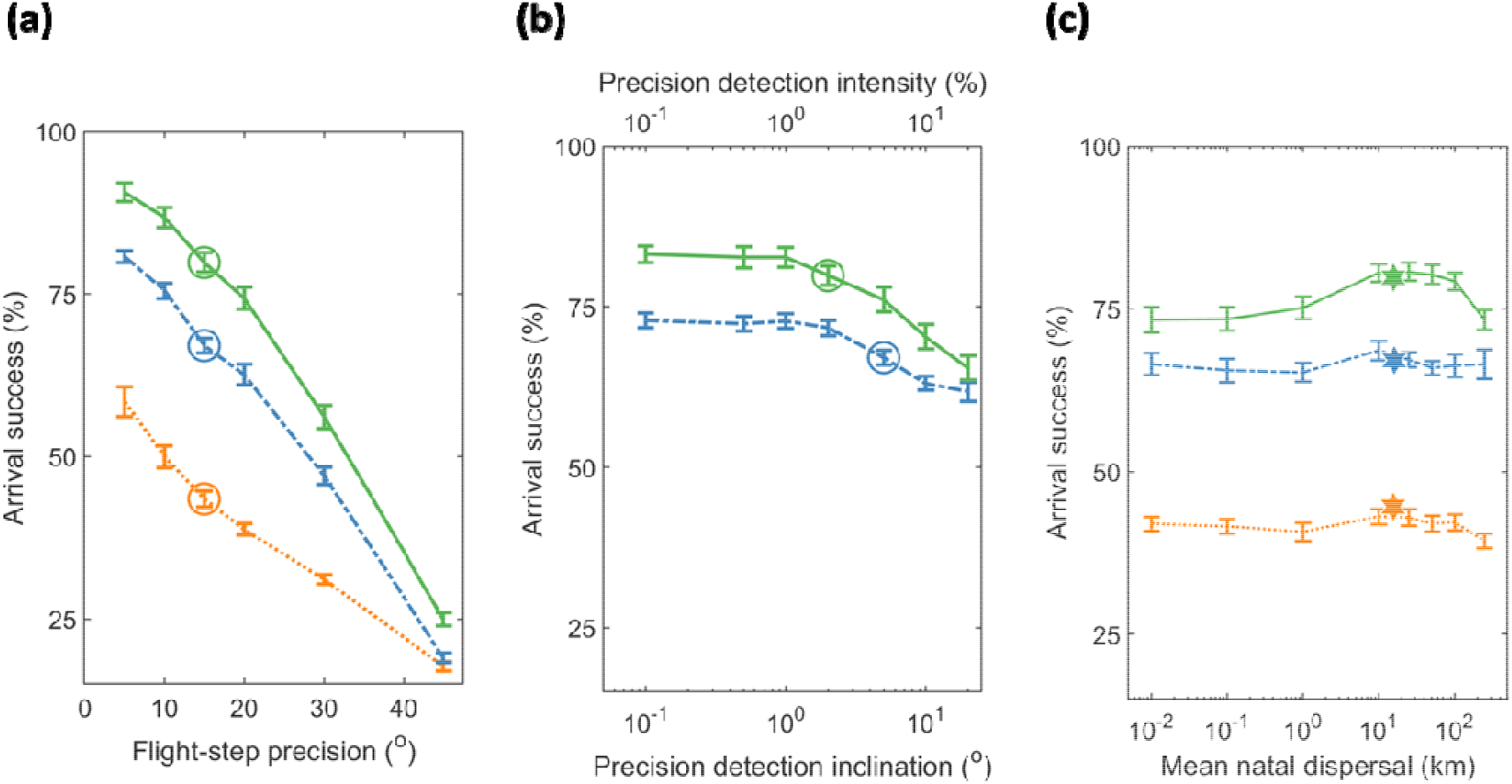
Robustness in arrival success of modelled magnetic-based migration to precision among flight-steps and in gauging signposts. All panels depict long-term mean arrival success (%) and standard deviation among years for non-signposted migration (orange dashed line), and signposted migration based on inclination (dot-dashed blue line) and intensity (solid green line). Arrival success is plotted as a function of (**a**) precision among flight-steps (degrees), (**b**) precision in gauging signposts based on inclination (degrees, dot-dashed blue line), and based on intensity (percent intensity, solid green line), and (**c**) non-evolved mean natal dispersal (i.e., distance between breeding and natal grounds, km). Circle symbols depict default parameters of (**a**)15° flight-flight-step precision and (**b**) 5° precision in detection of signpost inclination and 2% of intensity signpost. The hexagon symbols in (**c**) depict model-evolved mean natal dispersal (as in Fig. 3).

Overall, magnetic compass use was advantageous in comparison with geographic compass use. Table 2 compares, for default compass precision (circles in Fig. 4a-b), mean arrival success for all combinations of magnetic and geographic compass use, and for non-signposted migration, including with a declination signpost. Once again, the hierarchy in performance among non-signposted and signposted migration remained the same, with declination-signposted programs less successful compared with other signposted programs (52.4% arrival success with default all-magnetic compass use). Inheriting magnetic as opposed to geographic headings benefitted arrival success among signposted programs (median 15.9% relative gain in success, range 8.4 to 40.0%), but not so with non-signposted migration (median −0.6% relative gain, range −2.5 to 0.4%). Contrastingly, non-signposted migration with a primary magnetic compass always performed better (median 47.6%, range 43.1 to 54.8%) than with a primary star compass, but among signposted programs this effect was weaker (median 3.9%, range −1.9 to 31.8%). Finally, using a geographic (star) in-flight compass clearly increased performance of geographically-inherited programs with an inclination compass (median 15.0%, range 11.9 to 18.1%) but had no clear or consistent effect among other signposted programs (median 3.2%, range −3.3 to 13.6%), nor for non-signposted programs (median 0.8%, range −2.8 to 2.6%). Taken together, it is interesting that for both non-signposted and signposted programs, purely magnetic compass use (top row) consistently outperformed purely geographic compass use (bottom row; median 19.8%, range 9.3 to 42.2%).

**Table 2.** Arrival success for default and alternative (non-default) choice in compass and signpost use. Geometric-mean and between-year standard deviation in arrival success, with the inherited compass, primary migratory compass and in-flight compass can either be geomagnetic or geographic (star compass) based. Declination-signposted migration (last column) is also listed. The first row lists the default model, with all-magnetic compass use. The highest performance for each program (e.g., inclination-signposted) is listed in bold. In all cases, other parameters were set to default values (Table 1).

| Inherited headings | Primary compass | In-flight compass | Non-signposted | Inclination-signposted | Intensity-signposted | Declination-signposted |
| --- | --- | --- | --- | --- | --- | --- |
|  |  |  | Total | Total | Total | Total |
| Magnetic | Magnetic | Magnetic | 43.5±1.2% | 67.1±1.1% | 79.8±1.5% | 52.4±2.3 % |
|  |  | Geographic (star) | 43.5±0.6% | 70.2±1.9% | **82.2±2.3%** | **52.7±1.7%** |
|  | Geographic (star) | Magnetic | 28.9±0.5% | 68.4±2.9% | 78.8±2.7% | 43.7±3.3% |
|  |  | Geographic (star) | 28.1±0.3% | **70.9±3.6%** | 80.6±3.8% | 46.0±3.3% |
| Geographic | Magnetic | Magnetic | 43.1±1.3% | 53.2±3.4% | 72.9±1.6% | 47.7±2.1% |
|  |  | Geographic (star) | **43.8±1.1%** | 60.4±2.1% | 70.6±1.3% | 48.6±2.6% |
|  | Geographic (star) | Magnetic | 29.8±0.4% | 50.3±1.7% | 64.7±1.0% | 36.2±2.1% |
|  |  | Geographic (star) | 30.6±0.2% | 61.4±1.3% | 67.8±0.9% | 41.9±0.9% |

Both signposted programs evolved a sharp migratory divide in trans-Atlantic routes. Figure 5a-b illustrates the evolution of headings and *Zugknick* latitudes for intensity-signposted migration. Individuals breeding in NE and N Greenland evolved close to magnetic S headings (~180°) with *Zugknicks* in Africa, whereas individuals breeding in Canada and S and W Greenland, evolved closer to magnetic SW headings (~135°) with *Zugknicks* in W Europe. Inclination-signposted migration evolved a similarly SW-NE contrast in headings, but with individuals breeding in Quebec and the Southern tip of Greenland evolving more direct (but less successful) routes towards Africa rather than via Europe (Suppl. Fig 3). For intensity-signposted migrants, initial magnetic headings shifted clockwise (Fig. 5c, mean 16.5°) between 1900-2023, resulting in a Southward latitudinal shift in *Zugknicks* (Fig. 5d, mean 5.2°). Shifts in routes and their effect on arrival success varied over time and regionally: Suppl. Fig. 4 illustrates in detail for migration from the Western fringe of the population on Baffin Island (60°-80°W, 62.5°-70°N), how orientation shifts accelerated in the second half of the study period (1961-2023), resulting in enhanced (~10%) over-water mortality.

**Figure 5:**
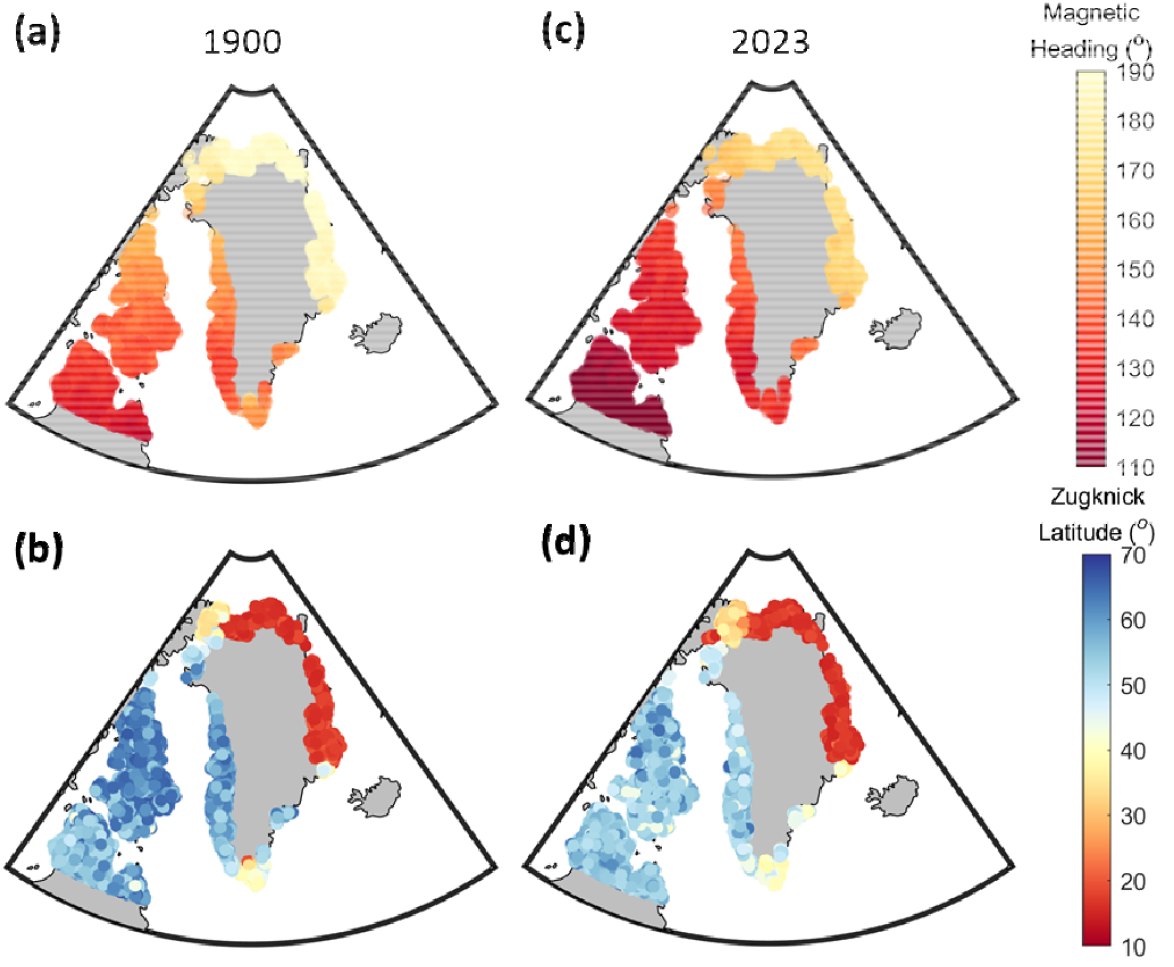
Evolution of modelled intensity-signposted migration of *leucoroha* wheatears to longterm geomagnetic shifts. Coloured symbols of 5000 randomly-selected successful modelled individuals from 1900 (**a-b**) and 2023 (**c-d**) illustrate (**a, c**) inherited magnetic headings (clockwise degrees from magnetic N) and (**b, d**) *Zugknick* latitudes (degrees). Stereographic azimuthal projection.

Robustness of magnetic-based migration to the strong geomagnetic changes was contingent on intrinsic variability in inheritance, as illustrated in Fig. 6 for intensity-signposted migration. Model-evolved standard deviations in inheritance of headings were consistent across migratory orientation programs, for example ranging from 2.3°-2.7° among non-signposted and non-signposted programs with default parameters, including across the range of tested distributions of natal dispersal (Fig. 3c), and also with smaller population sizes (Suppl. Fig. 3b). Model-evolved standard deviations in signposts were 0.58° for default inclination-signposted and 0.53% for default intensity-signposted programs, and in the sensitivity analysis were similar across all tested distributions of natal dispersal (0.54±0.2° and 0.54±0.03%). For the default intensity-signposted program, Fig. 6a depicts population-mean changes in inherited magnetic headings (circle colours) and intensity signposts (triangle colours), resulting in consistent arrival success (~80%) and limited over-water mortality (<25%). However, with perfect inheritance (zero standard deviations, Fig. 6b), headings failed to adapt while signposts increased, as over-water mortality became unsustainable (>50%). With geographic-inherited headings and compass use (Fig. 6c-d), arrival was overall lower but less dramatically regarding shifts in *Zugknick* locations, even without intrinsic variability in inheritance (Fig. 6d).

**Fig. 6:**
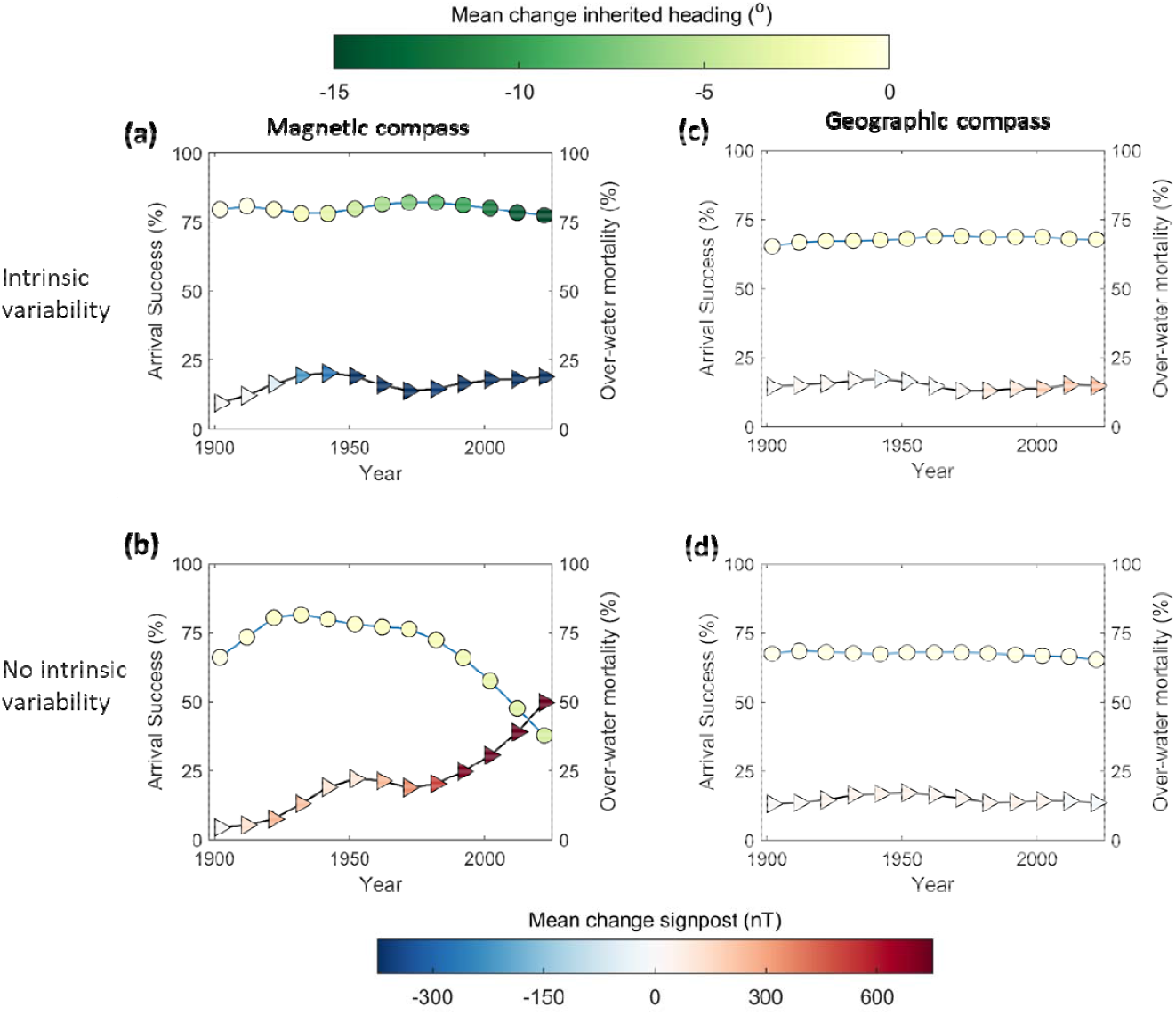
Intrinsic variability in inheritance facilitates evolution of successful magnetic headings and signposts through an extreme geomagnetic shift. Circles represent mean successful arrival (left axes) and triangles mortality over water (right axes) among intensity-signposted migrants (Fig. 2c), with symbol colours depicting population-mean changes in inherited headings since 1900 (circles, clockwise degrees from magnetic N) and in signpost magnitude (triangles, nT). (**a**) With model-evolved standard deviation in inherited headings (2.6°) and intensity signposts (0.53%),; (**b**) as (a) but without intrinsic standard deviations in inheritance of headings and signposts; (**c**) with an inherited geographic (star) primary and in-flight compass; (**d**) as in (**c**), but in the absence of intrinsic variability.

## Discussion

This study represents a first assessment of how migratory populations can adapt to complex and long-term shifts in geomagnetic landscapes, through natural selection of inherited magnetic information, together with a primary magnetic migratory compass. In particular, model results support the idea that inexperienced migrants can negotiate detoured routes using inherited magnetic headings and signposts. Such a gauge-and-compass program, where a magnetic “gauge” rather than circannual clock triggers switches in compass headings, could be particularly important for populations where naïve migrants travel independently, for example in providing a mechanism to evolve novel routes shaped by shifting habitat suitability and climate refugia (23,75,76). Magnetic signposts can in principle also trigger orientation shifts between celestial compass headings, such as a sunset compass or star compass (15,16). Depending on inner clock-updates and sensitivity to variable scheduling, sun-compass headings may be particularly advantageous over long-distance and high-latitude routes (16). Lastly, our results support the idea that intrinsic variability in migratory orientation, in addition to facilitating expansion of breeding and non-breeding ranges (11,77,78), is important in maintaining or modulating routes in unpredictable environments (36). While arrival success decreased slightly (~3%) in the rapidly-shifting final decades (Fig. 6a), this would have become prohibitive without intrinsic variability in inheritance (>40%; Fig. 6b). The extent to which such intrinsic variability would either be evolvable through natural selection or constrained by molecular (genetic or developmental) processes remains unknown.

For *leucorhoa* wheatears, it is unsurprising that *Zugknicks* via Europe are of adaptive benefit, given the significant risk of insufficient energy reserves for the minimally 4000-km direct open-ocean flights to Africa (20,51). Nonetheless, the persistent success of modelled magnetic-based migration (Figs. 3, 6a) across the rapidly-shifting geographic North pole region is somewhat remarkable, and illustrates that even strongly varying geomagnetic landscapes can be potentially advantageous relative to geographic-based movement (cf. 35) rather than necessarily represent a hazard (cf. 34). The fact that inherited magnetic headings and a primary magnetic compass generally outperformed their geographic counterparts (Table 2, Fig. 6) relates to the clockwise shift in declination (Fig. 1) which both reduces the ocean crossing (15,35) and facilitates self-correction in orientation (Suppl. Fig. 1). These factors are of course route-specific, so will affect the relative favourability of a primary magnetic compass (35) including its robustness to imprecision (16). It is further interesting that both inclination-signposted and intensity-signposted modelled populations evolved sharp SW-NE migratory divides, with the NE subpopulation (NE Greenland) almost not requiring a *Zugknick* to reach West Africa (Fig. 5, Suppl. Fig. 3). Such a scenario could lead to effective reproductive isolation through hybrid mortality (39,79), or alternatively development of dominance inheritance patterns between competing alleles (38), in contrast to the co-dominance (averaging) of traits modelled here. Naturally, the actual routes taken by *leucoroha* wheatears and their robustness to geomagnetic shifts could be modulated by other factors not explicitly considered, such as capacity for migratory endurance flight (51,80), reliability of selected winds (19,81) and avoidance of hazards associated with the longer detour (e.g., diminishing seasonal resources, exposure to predation (20)). Moreover, favourability among potential signposts will depend on the exact nature of the avian magnetic compass mechanism, which remains uncertain. With the favoured radical-pair magnetoreception, geomagnetic intensity will amplify the received signal (82), and the avian magnetic compass is known to require a discernible (though not necessarily precisely measurable) inclination angle (6,10,71). Therefore, it seems reasonable to conjecture that inexperienced migrants might be able to “gauge” a magnetic signal based on intensity, inclination or some combination of both (82,83). Gauging geomagnetic declination seems less likely for naïve migrants, since it requires comparison of geographic and geomagnetic axes, often while on the move and close to dark (16,84). Declination signposts additionally underperformed for modelled *leucoroha* wheatears, and were not sufficient for experienced migrants to correct for displacement in a recent experiment (8).

It is important to consider why inexperienced migrants might use a magnetic signpost to mediate detours rather than clearly important migratory cues such as habitat quality (22) or topography (62,85). However, such extrinsic cues may not always be reliably precise or unambiguous in negotiating long-distance routes. For example, an inherited *Zugknick* to reorient to the South on completing an open-ocean endurance flight could work for *leucoroha* wheatears arriving to Western Europe but, if they first stopped in Iceland, additional information would be required to avoid misorientation into the mid-Atlantic Ocean. We therefore propose that migratory orientation responses to coastal and habitat cues could be both inherited and mediated or triggered by magnetic information during the long-distance phase of migration, similarly to energetic and stopover decisions (86,87). Given that geomagnetic and celestial cues are fairly stable within migratory periods, we speculate that inherited compass information is typically primary, at least among long-distance nocturnally-migrating birds (6,16), though the degrees to which other environmental and social cues refine and modulate this programme remain to be clarified (4,88,89).

As an alternative to compass-based inherited orientation, it has also been proposed that naïve migrants might be able to perform navigation by following gradients in the geomagnetic field (9). The relative feasibility and efficiency of constant-heading vs. gradient-based migration remains an open question (9,16,32). For actual migration systems, this has only been assessed for migration based on correlated random walks – per definition less directed than compass courses – with supplemental navigational abilities based on geomagnetic information (32). While an innate or early-learned navigational ability offers the possibility to correct for imprecision or displacement, e.g., by currents, it could also produce inefficient migrations whenever gradients in field components are closely aligned (29,90). Our results suggest that constant-heading migration modulated by magnetic signposts could be sufficiently robust to variable and changing geomagnetic fields, at least given suitable intrinsic variability in inherited headings.

## Conclusions

While global patterns of avian migration can be explained as efficient energy acquisition of seasonal resources (91), enabling hindcasts of prehistoric migration routes (23,76), little is understood regarding the population consequences of how migratory orientation is transmitted across generations. Using an evolutionary algorithm approach enables population-level assessments of how inherited migratory orientation programs can both mediate and constrain adaptation of historic and novel migration routes. Our methods can be extended to assess other geophysical cues (e.g., sun azimuth) and flexible orientation reactions to other environmental factors such as currents, coastlines and habitat quality (24,54,85), including to assess resilience to climate change. More generally, our results illustrate how the Earth’s magnetic field may possibly play a vital role in the evolution of migration routes, as mediator between proximate environmental cues and ultimate drivers of population fitness through migratory success.

## Abbreviations

N: North
S: South
E: East
W: West

## Author contributions

All authors worked to conceive the study. J.D.M. formulated and coded the models, analysed the results, and wrote the original draft manuscript. H.S. and B.B. supervised the study and project, and contributed to manuscript revisions.

## Consent for publication

Not applicable.

## Availability of data and materials

The code to simulate the model and reproduce all the results figures is available in github repository (to be uploaded)

## References

1. Dingle H, Drake VA. What Is Migration? BioScience. 2007 Feb 1;57(2):113–21.

2. Alerstam T, Hedenström A, Åkesson S. Long-distance migration: evolution and determinants. Oikos. 2003;103(2):247–60.

3. Berdahl AM, Kao AB, Flack A, Westley PAH, Codling EA, Couzin ID, et al. Collective animal navigation and migratory culture: from theoretical models to empirical evidence. Phil Trans R Soc B. 2018 May 19;373(1746):20170009.

4. Aikens EO, Bontekoe ID, Blumenstiel L, Schlicksupp A, Flack A. Viewing animal migration through a social lens. Trends in Ecology & Evolution. 2022 Nov 1;37(11):985–96.

5. Bruderer B. The Study of Bird Migration by RadarPart 2: Major Achievements. Naturwissenschaften. 1997 Feb 1;84(2):45–54.

6. Mouritsen H. Long-distance navigation and magnetoreception in migratory animals. Nature. 2018 Jun;558(7708):50–9.

7. Lohmann KJ, Lohmann CMF. There and back again: natal homing by magnetic navigation in sea turtles and salmon. el Jundi B, Kelber A, Webb B, editors. Journal of Experimental Biology. 2019 Feb 6;222(Suppl_1):jeb184077.

8. Kishkinev D, Packmor F, Zechmeister T, Winkler HC, Chernetsov N, Mouritsen H, et al. Navigation by extrapolation of geomagnetic cues in a migratory songbird. Current Biology. 2021 Apr;31(7):1563–1569.e4.

9. Lohmann KJ, Goforth KM, Mackiewicz AG, Lim DS, Lohmann CMF. Magnetic maps in animal navigation. J Comp Physiol A. 2022 Jan 1;208(1):41–67.

10. Wiltschko R, Wiltschko W. Avian Navigation: A Combination of Innate and Learned Mechanisms. In: Advances in the Study of Behavior [Internet]. Elsevier; 2015 [cited 2021 Feb 5]. p. 229–310. Available from: https://linkinghub.elsevier.com/retrieve/pii/S0065345414000047

11. Berthold P. Control of Bird Migration. Springer Science & Business Media; 1996. 370 p.

12. Åkesson S, Helm B. Endogenous Programs and Flexibility in Bird Migration. Frontiers in Ecology and Evolution [Internet]. 2020 [cited 2023 Feb 12];8. Available from: https://www.frontiersin.org/articles/10.3389/fevo.2020.00078

13. Åkesson S, Ilieva M, Karagicheva J, Rakhimberdiev E, Tomotani B, Helm B. Timing avian long-distance migration: from internal clock mechanisms to global flights. Philosophical Transactions of the Royal Society B: Biological Sciences. 2017 Nov 19;372(1734):20160252.

14. Åkesson S, Bianco G. Route simulations, compass mechanisms and long-distance migration flights in birds. J Comp Physiol A. 2017 Jul;203(6–7):475–90.

15. Muheim R, Schmaljohann H, Alerstam T. Feasibility of sun and magnetic compass mechanisms in avian long-distance migration. Mov Ecol. 2018 Dec;6(1):8.

16. McLaren JD, Schmaljohann H, Blasius B. Predicting performance of naïve migratory animals, from many wrongs to self-correction. Commun Biol. 2022 Oct 4;5(1):1–16.

17. Thorup K, Rabøl J, Erni B. Estimating variation among individuals in migration direction. Journal of Avian Biology. 2007 Mar;38(2):182–9.

18. Erni B, Liechti F, Bruderer B. The role of wind in passerine autumn migration between Europe and Africa. Behavioral Ecology. 2005 Jul 1;16(4):732–40.

19. McLaren JD, Shamoun-Baranes J, Bouten W. Wind selectivity and partial compensation for wind drift among nocturnally migrating passerines. Behavioral Ecology. 2012;23(5):1089–101.

20. Alerstam T. Detours in Bird Migration. Journal of Theoretical Biology. 2001 Apr 7;209(3):319–31.

21. Cohen EB, Barrow WC, Buler JJ, Deppe JL, Farnsworth A, Marra PP, et al. How do en route events around the Gulf of Mexico influence migratory landbird populations? cond. 2017 May;119(2):327–43.

22. Thorup K, Tøttrup AP, Willemoes M, Klaassen RHG, Strandberg R, Vega ML, et al. Resource tracking within and across continents in long-distance bird migrants. Science Advances. 2017 Jan 4;3(1):e1601360.

23. Thorup K, Pedersen L, da Fonseca RR, Naimi B, Nogués-Bravo D, Krapp M, et al. Response of an Afro-Palearctic bird migrant to glaciation cycles. Proc Natl Acad Sci USA. 2021 Dec 28;118(52):e2023836118.

24. Shamoun-Baranes J, Liechti F, Vansteelant WMG. Atmospheric conditions create freeways, detours and tailbacks for migrating birds. J Comp Physiol A. 2017 Jul;203(6–7):509–29.

25. Gill JA, Alves JA, Gunnarsson TG. Mechanisms driving phenological and range change in migratory species. Phil Trans R Soc B. 2019 Sep 16;374(1781):20180047.

26. Gwinner E, Wiltschko W. Endogenously controlled changes in migratory direction of the garden warbler,Sylvia borin. J Comp Physiol. 1978 Sep 1;125(3):267–73.

27. Laundal KM, Richmond AD. Magnetic Coordinate Systems. Space Sci Rev. 2017 Mar;206(1–4):27–59.

28. Rakhimberdiev E, Karagicheva J, Jaatinen K, Winkler DW, Phillips JB, Piersma T. Naïve migrants and the use of magnetic cues: temporal fluctuations in the geomagnetic field differentially affect male and female Ruff *Philomachus pugnax* during their first migration. Bauer S, editor. Ibis. 2014 Oct;156(4):864–9.

29. Pizzuti S, Bernish M, Harvey A, Tourangeau L, Shriver C, Kehl C, et al. Uncovering how animals use combinations of magnetic field properties to navigate: a computational approach. J Comp Physiol A. 2022 Jan 1;208(1):155–66.

30. Hongre L, Sailhac P, Alexandrescu M, Dubois J. Nonlinear and multifractal approaches of the geomagnetic field. Physics of the Earth and Planetary Interiors. 1999 Feb;110(3–4):157–90.

31. Livermore PW, Finlay CC, Bayliff M. Recent north magnetic pole acceleration towards Siberia caused by flux lobe elongation. Nat Geosci. 2020 May;13(5):387–91.

32. Zein B, Long JA, Safi K, Kölzsch A, Wikelski M, Kruckenberg H, et al. Simulation experiment to test strategies of geomagnetic navigation during long-distance bird migration. Movement Ecology. 2021 Sep 15;9(1):46.

33. Sjöberg S, Muheim R. A New View on an Old Debate: Type of Cue-Conflict Manipulation and Availability of Stars Can Explain the Discrepancies between Cue-Calibration Experiments with Migratory Songbirds. Front Behav Neurosci [Internet]. 2016 Feb 23 [cited 2021 Feb 5];10. Available from: http://journal.frontiersin.org/Article/10.3389/fhbeh.2016.00029/abstract

34. Kok EMA, Tibbitts TL, Douglas DC, Howey PW, Dekinga A, Gnep B, et al. A red knot as a black swan: how a single bird shows navigational abilities during repeat crossings of the Greenland Icecap. Journal of Avian Biology [Internet]. 2020 [cited 2022 Jun 30];51(8). Available from: https://onlinelibrary.wiley.com/doi/abs/10.1111/jav.02464

35. Alerstam T. Evaluation of Long-Distance Orientation in Birds on the Basis of Migration Routes Recorded by Radar and Satellite Tracking. J Navigation. 2001 Sep;54(3):393–403.

36. Reilly JR, Reilly RJ. Bet-hedging and the orientation of juvenile passerines in fall migration: Bet-hedging in passerine migration. Journal of Animal Ecology. 2009 Jul 29;78(5):990–1001.

37. Delmore K, Illera JC, Pérez-Tris J, Segelbacher G, Lugo Ramos JS, Durieux G, et al. The evolutionary history and genomics of European blackcap migration. Scordato E, Wittkopp PJ, editors. eLife. 2020 Apr 21;9:e54462.

38. Sokolovskis K, Lundberg M, Åkesson S, Willemoes M, Zhao T, Caballero-Lopez V, et al. Migration direction in a songbird explained by two loci. Nat Commun. 2023 Jan 11;14(1):165.

39. Delmore KE, Fox JW, Irwin DE. Dramatic intraspecific differences in migratory routes, stopover sites and wintering areas, revealed using light-level geolocators. Proceedings of the Royal Society B: Biological Sciences. 2012 Nov 22;279(1747):4582–9.

40. Weatherhead PJ, Forbes MRL. Natal philopatry in passerine birds: genetic or ecological influences? Behavioral Ecology. 1994 Dec 1;5(4):426–33.

41. Hansson B, Bensch S, Hasselquist D. Heritability of dispersal in the great reed warbler. Ecology Letters. 2003;6(4):290–4.

42. Simons AM. The continuity of microevolution and macroevolution. Journal of Evolutionary Biology. 2002;15(5):688–701.

43. Rudolph G. Evolutionary Strategies. In: Rozenberg G, Bäck T, Kok JN, editors. Handbook of Natural Computing [Internet]. Berlin, Heidelberg: Springer; 2012 [cited 2022 Aug 4]. p. 673–98. Available from: https://doi.org/10.1007/978-3-540-92910-9_22

44. Bairlein F, Norris DR, Nagel R, Bulte M, Voigt CC, Fox JW, et al. Cross-hemisphere migration of a 25 g songbird. Biol Lett. 2012 Aug 23;8(4):505–7.

45. Boone RB. Evolutionary computation in zoology and ecology. Current Zoology. 2017 Dec 1;63(6):675–86.

46. De Jong K. Generalized Evolutionary Algorithms. In: Rozenberg G, Bäck T, Kok JN, editors. Handbook of Natural Computing [Internet]. Berlin, Heidelberg: Springer; 2012 [cited 2022 Aug 4]. p. 625–35. Available from: https://doi.org/10.1007/978-3-540-92910-9_20

47. Wagner GP, Altenberg L. Perspective: Complex Adaptations and the Evolution of Evolvability. Evolution. 1996;50(3):967–76.

48. Thébault E, Finlay CC, Beggan CD, Alken P, Aubert J, Barrois O, et al. International Geomagnetic Reference Field: the 12th generation. Earth Planet Sp. 2015 Dec;67(1):79.

49. Chernetsov N. Compass systems. J Comp Physiol A. 2017 Jul;203(6–7):447–53.

50. Komolkin AV, Kupriyanov P, Chudin A, Bojarinova J, Kavokin K, Chernetsov N. Theoretically possible spatial accuracy of geomagnetic maps used by migrating animals. J R Soc Interface. 2017 Mar;14(128):20161002.

51. Bulte M, McLaren JD, Bairlein F, Bouten W, Schmaljohann H, Shamoun-Baranes J. Can wheatears weather the Atlantic? Modeling nonstop trans-Atlantic flights of a small migratory songbird. The Auk. 2014 Jul;131(3):363–70.

52. Thorup K, Ortvad TE, Rabøl J. Do Nearctic Northern Wheatears (Oenanthe oenanthe leuchorhoa) migrate nonstop to Africa? :6.

53. Ottosson U, Sandberg R, Pettersson J. Orientation Cage and Release Experiments with Migratory Wheatears (Oenanthe oenanthe) in Scandinavia and Greenland: The Importance of Visual Cues. Ethology. 1990;86(1):57–70.

54. Schmaljohann H, Naef-Daenzer B. Body condition and wind support initiate the shift of migratory direction and timing of nocturnal departure in a songbird: Departure behaviour of free-flying birds. Journal of Animal Ecology. 2011 Nov;80(6):1115–22.

55. Schmaljohann H, Meier C, Arlt D, Bairlein F, van Oosten H, Morbey YE, et al. Proximate causes of avian protandry differ between subspecies with contrasting migration challenges. Behavioral Ecology. 2016 Jan 1;27(1):321–31.

56. Cramp S (chief ed). Handbook of Europe the Middle East and North Africa, Volume 5: Tyrant Flycatchers to Thrushes [Internet]. Oxford: Oxford University Press; 1988 [cited 2022 Jun 30]. Available from: https://www.biblio.com/book/handbook-europe-middle-east-north-africa/d/1336501969

57. Mardia KV. Statistics of Directional Data. Journal of the Royal Statistical Society Series B (Methodological). 1975;37(3):349–93.

58. Emmerich M, Shir OM, Wang H. Evolution Strategies. In: Martí R, Panos P, Resende MGC, editors. Handbook of Heuristics [Internet]. Cham: Springer International Publishing; 2018 [cited 2022 Aug 4]. p. 1–31. Available from: https://doi.org/10.1007/978-3-319-07153-4_13-1

59. Ceresa F, Belda EJ, Monrós JS. Similar dispersal patterns between two closely related birds with contrasting migration strategies. Population Ecology. 2016;58(3):421–7.

60. Hu T, Banzhaf W. Evolvability and Speed of Evolutionary Algorithms in Light of Recent Developments in Biology. Journal of Artificial Evolution and Applications. 2010 Jun 2;2010:e568375.

61. Thornton PE, Rosenbloom NA. Ecosystem model spin-up: Estimating steady state conditions in a coupled terrestrial carbon and nitrogen cycle model. Ecological Modelling. 2005 Nov 25;189(1):25–48.

62. Erni B, Liechti F, Bruderer B. How Does a First Year Passerine Migrant Find Its Way? Simulating Migration Mechanisms and Behavioural Adaptations. Oikos. 2003;103(2):333–40.

63. Simons AM. Modes of response to environmental change and the elusive empirical evidence for bet hedging. Proceedings of the Royal Society B: Biological Sciences. 2011 Jun 7;278(1712):1601–9.

64. McLaren JD, Shamoun-Baranes J, Bouten W. Stop early to travel fast: modelling risk-averse scheduling among nocturnally migrating birds. Journal of Theoretical Biology. 2013 Jan;316:90–8.

65. Schmaljohann H, Eikenaar C, Sapir N. Understanding the ecological and evolutionary function of stopover in migrating birds. Biological Reviews [Internet]. [cited 2022 Jun 30];n/a(n/a). Available from: https://onlinelibrary.wiley.com/doi/abs/10.1111/brv.12839

66. Tuanmu MN, Jetz W. A global 1-km consensus land-cover product for biodiversity and ecosystem modelling. Global Ecology and Biogeography. 2014;23(9):1031–45.

67. Müller F, Taylor PD, Sjöberg S, Muheim R, Tsvey A, Mackenzie SA, et al. Towards a conceptual framework for explaining variation in nocturnal departure time of songbird migrants. Movement Ecology. 2016 Oct 17;4(1):24.

68. Jenkins A. The Sun’s position in the sky. Eur J Phys. 2013 May 1;34(3):633–52.

69. Liechti F. Nächtlicher Vogelzug im Herbst über Süddeutschland: Winddrift und Kompensation. J Ornithol. 1993 Oct 1;134(4):373–404.

70. Bäckman J, Alerstam T. Orientation scatter of free-flying nocturnal passerine migrants: components and causes. Animal Behaviour. 2003 May;65(5):987–96.

71. Lefeldt N, Dreyer D, Schneider NL, Steenken F, Mouritsen H. Migratory blackcaps tested in Emlen funnels can orient at 85 degrees but not at 88 degrees magnetic inclination. Journal of Experimental Biology. 2015 Jan 15;218(2):206–11.

72. Greene CA, Thirumalai K, Kearney KA, Delgado JM, Schwanghart W, Wolfenbarger NS, et al. The Climate Data Toolbox for MATLAB. Geochemistry, Geophysics, Geosystems. 2019;20(7):3774–81.

73. Compston D. International Geomagnetic Reference Field (IGRF) Model [Internet]. Available from: https://www.mathworks.com/matlabcentral/fileexchange/34388-international-geomagnetic-reference-field-igrf-model

74. Brown W. 13th Generation International Geomagnetic Reference Field [Internet]. Available from: https://github.com/wb-bgs/m_IGRF

75. Zink RM, Gardner AS. Glaciation as a migratory switch. Science Advances. 2017 Sep 20;3(9):e1603133.

76. Somveille M, Wikelski M, Beyer RM, Rodrigues ASL, Manica A, Jetz W. Simulation-based reconstruction of global bird migration over the past 50,000 years. Nat Commun. 2020 Feb 18;11(1):801.

77. Veit RR, Manne LL, Zawadzki LC, Alamo MA, Henry RW. Editorial: Vagrancy, exploratory behavior and colonization by birds: Escape from extinction? Frontiers in Ecology and Evolution [Internet]. 2022 [cited 2022 Dec 5];10. Available from: https://www.frontiersin.org/articles/10.3389/fevo.2022.960841

78. Dufour P, Åkesson S, Hellström M, Hewson C, Lagerveld S, Mitchell L, et al. The Yellow-browed Warbler (Phylloscopus inornatus) as a model to understand vagrancy and its potential for the evolution of new migration routes. Movement Ecology. 2022 Dec 14;10(1):59.

79. Hendry AP.* Selection against migrants contributes to the rapid evolution of ecologically dependent reproductive isolation. Evol Ecol Res. 2004;6(8):1219–36.

80. Delingat J, Bairlein F, Hedenström A. Obligatory barrier crossing and adaptive fuel management in migratory birds: the case of the Atlantic crossing in Northern Wheatears (Oenanthe oenanthe). Behav Ecol Sociobiol. 2008 May;62(7):1069–78.

81. Schmaljohann H, Korner-Nievergelt F, Naef-Daenzer B, Nagel R, Maggini I, Bulte M, et al. Stopover optimization in a long-distance migrant: the role of fuel load and nocturnal take-off time in Alaskan northern wheatears (Oenanthe oenanthe). Front Zool. 2013;10(1):26.

82. Xu J, Jarocha LE, Zollitsch T, Konowalczyk M, Henbest KB, Richert S, et al. Magnetic sensitivity of cryptochrome 4 from a migratory songbird. Nature. 2021 Jun;594(7864):535–40.

83. Worster S, Mouritsen H, Hore PJ. A light-dependent magnetoreception mechanism insensitive to light intensity and polarization. J R Soc Interface. 2017 Sep;14(134):20170405.

84. Muheim R, Philips JB, Åkesson S. Polarized Light Cues Underlie Compass Calibration in Migratory Songbirds. Science. 2006 Aug 11;313(5788):837–9.

85. Aurbach A, Schmid B, Liechti F, Chokani N, Abhari R. Complex behaviour in complex terrain - Modelling bird migration in a high resolution wind field across mountainous terrain to simulate observed patterns. Journal of Theoretical Biology. 2018 Oct;454:126–38.

86. Kullberg C, Henshaw I, Jakobsson S, Johansson P, Fransson T. Fuelling decisions in migratory birds: geomagnetic cues override the seasonal effect. Proceedings of the Royal Society B: Biological Sciences. 2007 Jun 26;274(1622):2145–51.

87. Fransson T, Jakobsson S, Johansson P, Kullberg C, Lind J, Vallin A. Magnetic cues trigger extensive refuelling. Nature. 2001 Nov;414(6859):35–6.

88. Flack A, Aikens EO, Kölzsch A, Nourani E, Snell KRS, Fiedler W, et al. New frontiers in bird migration research. Current Biology. 2022 Oct 24;32(20):R1187–99.

89. Piersma T, Loonstra AHJ, Verhoeven MA, Oudman T. Rethinking classic starling displacement experiments: evidence for innate or for learned migratory directions? Journal of Avian Biology [Internet]. 2020 [cited 2022 Apr 24];51(5). Available from: https://onlinelibrary.wiley.com/doi/abs/10.1111/jav.02337

90. Boström JE, Åkesson S, Alerstam T. Where on earth can animals use a geomagnetic bicoordinate map for navigation? Ecography. 2012 Nov;35(11):1039–47.

91. Somveille M, Rodrigues ASL, Manica A. Energy efficiency drives the global seasonal distribution of birds. Nat Ecol Evol. 2018 Jun;2(6): 962–9.

