## Supplementary Material for "Gauge-and-compass migration: inherited magnetic headings and signposts can adapt to changing geomagnetic landscapes"

| 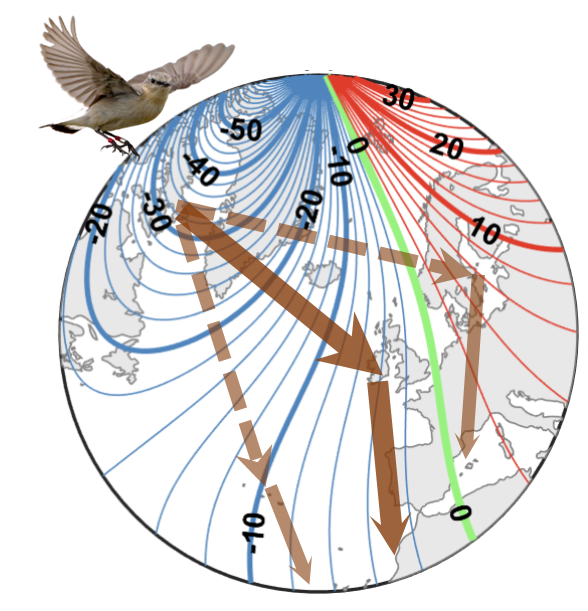 | **Suppl. Fig. 1**: Positive longitudinal gradients in geomagnetic declination facilitate self-correction using a geomagnetic compass. Contours of declination (clockwise degrees between true geographic and magnetic N) for 2010, together with likely routes taken by *leucoroha* wheatears (photo, HS). Solid thick brown arrows represent approximate route taken from Baffin Island, Canada and West Africa (Fig. 1, main text). If displaced *en route* (dashed brown arrows), the resultant *Zugknick* flight direction (thin solid brown arrows) is partially compensating, due to the counter-clockwise declination shift when displaced Westward, and clockwise shift when displaced Eastward. |
| --- | --- |

| 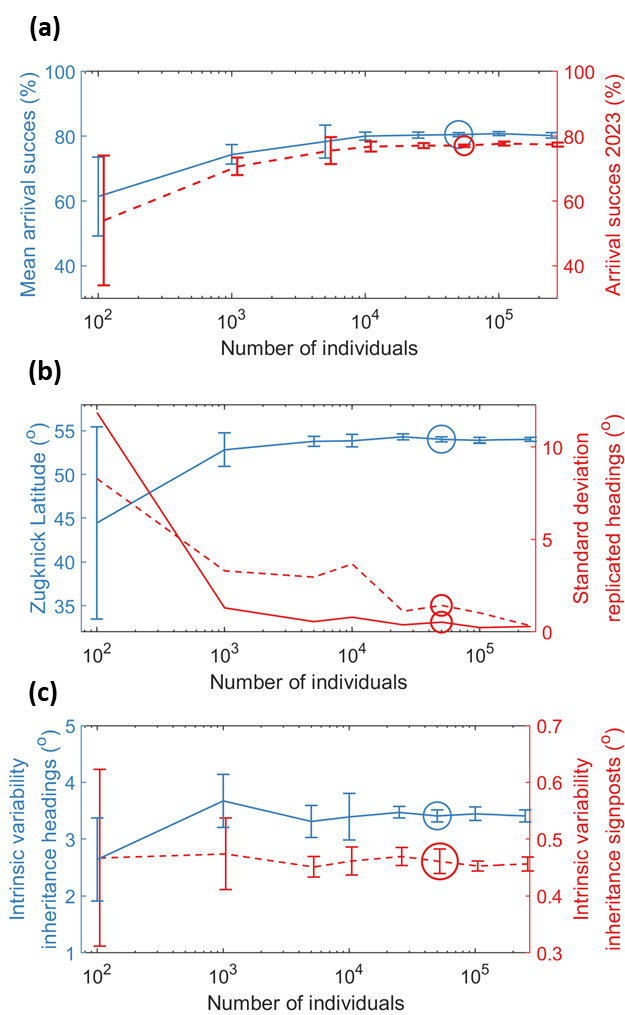 | **Suppl. Fig. 2**: Validation of model consistency with number of modelled individuals, using 6 replicates for intensity-signposted migration, (**a**) geometric-mean arrival success (blue lines) and final-year success (dashed red line) with, here and elsewhere, error bars depicting standard deviation over replicates; (**b**) population-mean final-year *Zugknick* latitude (blue lines and error bars) and between-replicate standard deviation in population-mean final-year inherited headings (red lines) and *Zugknick* headings (dashed red lines), all in degrees, (**c**) as evolved in the model spin-up, intrinsic standard deviation in inheritance of migratory headings (degrees, blue lines and error bars) and inheritance of intensity-signposts (% of actual field concentration, red lines and error bars). |
| --- | --- |

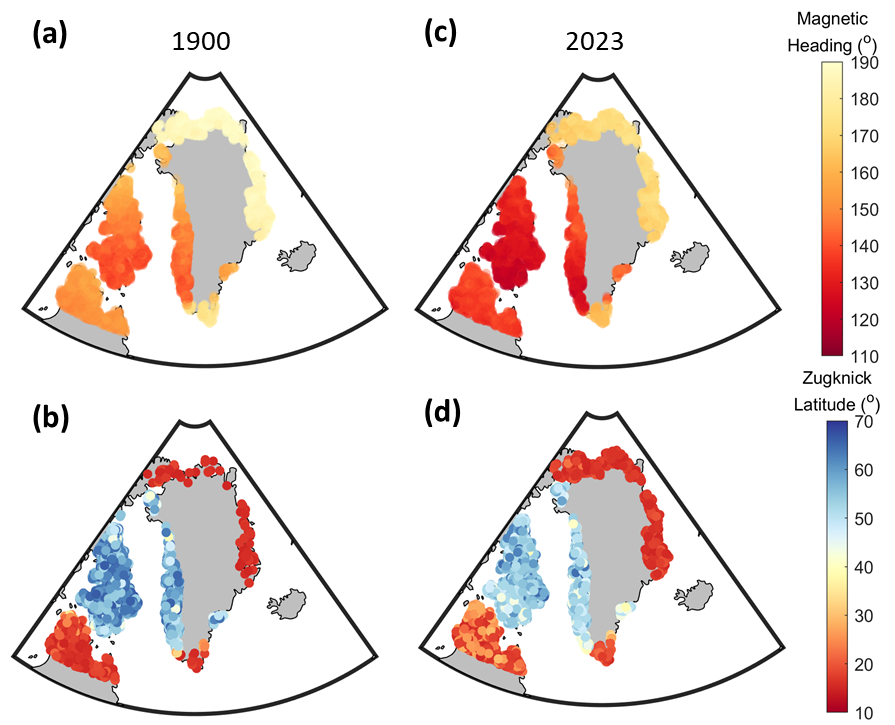

**Suppl. Fig. 3**: Evolution of modelled inclination-signposted migration of *leucoroha* wheatears to long-term geomagnetic shifts As in Fig. 5, but for inclination-signposted migration. Coloured symbols represent 5000 randomly-sampled inherited geomagnetic headings (**a**) and *Zugknick* latitude (**b**) in 1900, illustrate a SW/NE divide in headings but three-way division in migratory connectivity (individuals breeding in the NE and S/SW evolving direct routes to Africa, and individuals breeding in the W detoured routes. In 2023, (**c**) inherited magnetic headings shifted clockwise (mean 16.2°) while (**d**) *Zugknicks* underwent small Southward shifts (mean 3.8°). Stereographic azimuthal projection.

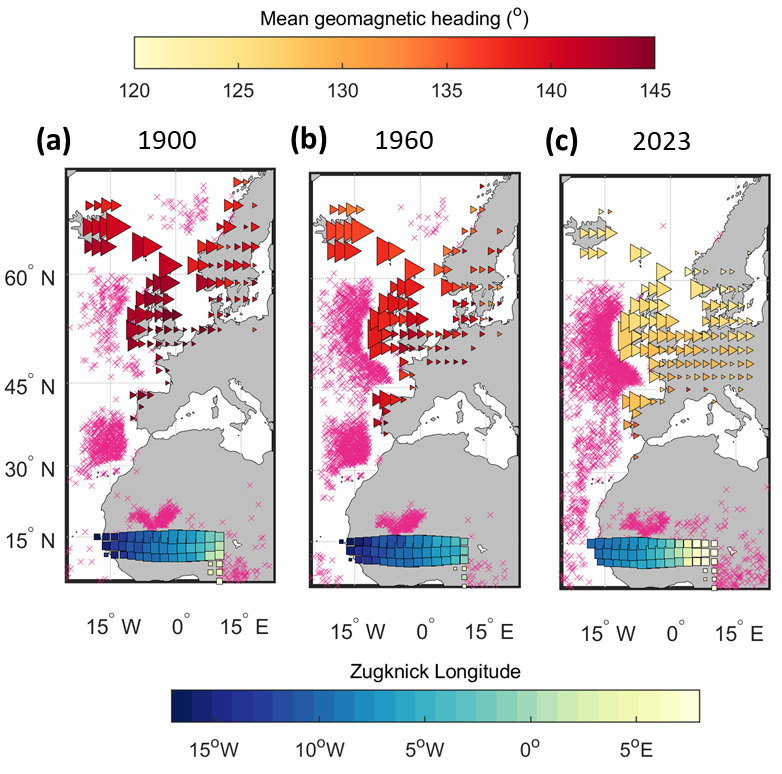

**Suppl. Fig. 4**: Increasing effects of modulated headings on routes and arrival of among modelled intensity-signposted *leucoroha* wheatears. Locations of *Zugknicks* (triangles) and arrival points at the wintering grounds (overlapping squares) of modelled *leucoroha* wheatear migration from Baffin Island (60°-80°W, 62.5°-70°N) in (**a**) 1900, (**b**) 1960 to (**c**) 2023. Symbol sizes are proportional to frequency of occurrence within 2°x2° areas, with triangle colours depicting mean inherited headings (clockwise degrees from magnetic N) and square colours mean *Zugknick* longitude. Pink crosses depict mortality over water, desert or having overshot the arrival area to the South or East.

In 1900, *Zugknicks* occurred most frequently between Iceland and Scotland (triangles in Suppl. Fig. 4a), following fairly uniform magnetic headings, with broad-front arrival to Africa (squares, coloured by *Zugknick* longitude with size scaled by frequency). By 1960 (Suppl. Fig. 4b), *Zugknicks* were shifted slightly towards Ireland (mean 2.2° Southward and 1.9° Westward shifts), with mean magnetic headings shifted 5.1° counter-clockwise (NW) but mean initial departure directions (geographic headings) shifted 8.6° clockwise (SE), resulting in 6.5% higher over-water mortality (pink crosses). By 2023 (Suppl. Fig. 4c), magnetic headings were shifted farther counter-clockwise (15.9°) yet initial geographic headings farther clockwise (23.5°), resulting in more continental *Zugknick* locations (mean 3.2° Eastward and 7.3° Southward shifts), and 9.8% higher over-water mortality compared with 1900.
